# Chlomito: a novel tool for precise elimination of organelle genome contamination in nuclear genome assemblies

**DOI:** 10.1101/2024.02.28.582616

**Authors:** Wei Song, Chong Li, Yanming Lu, Dawei Shen, Yunxiao Jia, Yixin Huo, Weilan Piao, Hua Jin

## Abstract

Accurate genome assemblies are crucial for understanding biological evolution, mechanisms of disease, and biodiversity. However, contamination from organelle genomes in nuclear genome analyses often leads to inaccuracies and unreliability in results. To address this issue, we developed a tool named Chlomito, which employs innovative algorithms to precisely identify and eliminate organelle genome contamination sequences from nuclear genome assemblies. Compared to conventional approaches, Chlomito can not only detect and eliminate organelle sequences but also effectively distinguish true organelle sequences from those transferred into the nucleus via horizontal gene transfer. To evaluate the accuracy of Chlomito, we conducted tests using sequencing data from Plum and Mango. The results confirmed that Chlomito can accurately detect contigs originating from the organelle genome, and the identified contigs covered most regions of the organelle reference genomes, demonstrating its efficiency and precision in comprehensively recognizing organelle genome sequences. Additionally, for user convenience, we packaged this method into a Docker image, simplifying the data processing workflow. Overall, Chlomito provides a highly efficient and accurate method for identifying and removing contigs derived from organelle genomes in genomic assembly data, thereby contributing to the improvement of genome assembly quality and advancing research in genomics and evolutionary biology.

## Introduction

With the widespread application of high-throughput sequencing technologies, researchers can rapidly obtain genomic sequences of various species (Rhoads and Au, 2015;Goodwin et al., 2016;Jain et al., 2018). However, during the process of genome assembly, the issue of contamination with organelle genomes often arises, where DNA sequences from mitochondria and chloroplasts may be mistakenly assembled into the nuclear genome. This phenomenon occurs because organelle DNA and nuclear DNA are extracted and sequenced simultaneously, resulting in mixed data sets. Additionally, since organelle genomes exist in multiple copies within the cell and are relatively small compared to the nuclear genome(Pyke, 1999), they are overrepresented in the sequencing data, which increases the risk of their erroneous assembled into the nuclear genome assembly.

Furthermore, sequences from organelle genomes may exhibit high similarity to certain sequences in the nuclear genome, for instance, organelle genes may be transferred into the nuclear genome due to horizontal gene transfer events(Martin, 2003;Timmis et al., 2004;Wei et al., 2022;Wang et al., 2024). This makes it challenging to accurately distinguish contigs derived from organelle genomes among all the assembled contigs, especially those containing sequences transferred from the organelle genome through horizontal gene transfer (HGT). Therefore, accurate identification and elimination of organelle genome sequences are essential for minimizing contamination issues, enhancing the quality and accuracy of genome assembly.

To identify organelle sequences from genome assembly data, the current widely adopted method relies on aligning the genome contigs to organelle reference genome and filtering based on alignment lengths (Howe et al., 2021;Mishra et al., 2021;Rhie et al., 2021;Zhang et al., 2024) or sequence similarity (Shirasawa et al., 2021;Bae et al., 2023;Yu et al., 2024). However, a significant limitation of this method is that it ignores the fact that organelle genomes can be inserted into the nuclear genome via horizontal gene transfer (HGT)(Cecchin et al., 2019;Allio et al., 2020;Kenny et al., 2020;Martin et al., 2023). Due to such transfers, fragments of the organelle genome may reside in the nuclear genome, making it challenging for traditional filtering methods to differentiate between the original organelle genome and the organelle DNA transferred into the nucleus via HGT. Additionally, existing methods often require access to pre-assembled organelle reference genomes, limiting applicability especially for species without well-assembled organelle references. Finally, the implementation of current methods generally lacks integrated and user-friendly software tool support, requiring researchers to manually perform all the steps, which is not only time-consuming but also prone to errors, particularly when dealing with large amounts of data.

To address the challenge of accurately identifying organelle genome sequences from genomic assemblies, we propose a novel solution in this study. Our approach specifically employs two core metrics: the alignment length coverage ratio(ALCR) and the sequencing depth ratio(SDR) to enhance the identification accuracy. The ALCR refers to the proportion of a contig’s length that aligns with the organelle reference genome relative to the total length of the contig. This metric can help differentiate contigs that contain only small fragments of organelle DNA, which may arise from horizontal gene transfer, as these fragments usually constitute only a small portion of the total contig length. Therefore, a low ALCR may indicate that the contig belongs to the nuclear genome rather than the organelle genome (Zhu et al., 2021;Nath et al., 2022;Hao et al., 2023;Zhou et al., 2023b). Meanwhile, the SDR refers to the ratio of each coting’s sequencing depth to the average sequencing depth of the organelle genome. Given that organelle genomes exist in multiple copies within a cell, they typically exhibit higher sequencing depths than nuclear genome(Sanita Lima et al., 2016;Wang et al., 2018;Li et al., 2021;Giorgashvili et al., 2022;Zhou et al., 2023a). Therefore, a contig with a high sequencing depth ratio, similar to the average of the organelle genome, is more likely to be a part of the organelle genome. By combining these two metrics, we can significantly improve the accuracy of identifying and removing organelle genome sequences from genome assembly data.

Furthermore, to facilitate usage by researchers with limited bioinformatics experience, we have implemented this new approach as easy-to-use software and packaged it as a Docker image, enabling easy distribution and execution across diverse computing platforms with a single command. Lastly, to validate the accuracy and reliability of our tool, we conducted tests using sequencing data from Plum (Prunus salicina) (Liu et al., 2020) and Mango (Mangifera indica) (Wang et al., 2020). The results demonstrate that our software can not only accurately identify organelle genome contigs from genome assemblies, but also accurately distinguish native organelle sequences from those inserted into the nuclear genome via horizontal gene transfer. In summary, our tool provides an accurate and effective solution for identifying and removing organelle DNA fragments from genome assembly contigs, holding significant value in improving the quality of chromosome assembly and deepening our understanding of the complex interactions between organelle and nuclear genomes.

## Material && Methods

### Availability of data and materials

To validate the accuracy of the Chlomito software in detecting organelle genome sequences, we utilized sequencing data of Mango and Plum from the NCBI Bioproject database, with accession numbers PRJNA487154 and PRJNA574159, respectively. These datasets include high-quality second and third-generation sequencing data, utilized for organelle genome identification and chromosome-level genome assembly.

### Implementation and installation of Chlomito

Chlomito is a Python-based software provided in the form of a Docker image. The image is accessible at https://hub.docker.com/repository/docker/songweidocker/chlomito. All experiments were conducted on an Ubuntu Linux 18.04.3 server, equipped with two Intel Xeon processors (32 cores each, totaling 64 threads) and 512 GB of RAM. Chlomito can be installed using the Docker command: docker pull songweidocker/chlomito:v1.

### Contig-level genome assembly

Flye (Kolmogorov et al., 2019) is a genome assembler software designed for long-read sequencing data from third-generation platforms such as PacBio and Oxford Nanopore. It is capable of assembling raw error-prone long reads into contiguous genomic sequences known as contigs. The goal of Flye is to generate high-quality genome assemblies, especially for large or complex genomes. In this study, we utilized Flye to assemble PacBio sequencing data of Mango and Plum into genome assemblies contigs. Due to the high error rate of third-generation sequencing data, the assembled contigs were then corrected using Racon (Vaser et al., 2017) and Pilon (Walker et al., 2014).

### Construction of organelle genome database

The organelle genomes, mitochondria and chloroplasts, were firstly assembled from second-generation sequencing data using GetOrganelle (Jin et al., 2020). GetOrganelle is a powerful genomics software tool specifically designed for efficient assembly of mitochondrial and chloroplast genomes. It is capable of simultaneously assembling organelle genomes from both mitochondria and chloroplasts. Compared to other similar software tools, GetOrganelle demonstrates superior performance in terms of both accuracy and speed for organelle genome assembly. After that, we have merged the mitochondrial and chloroplast genomes published in the NCBI organelle database with organelle genomes assembled using the GetOrganelle software, creating a comprehensive local organelle genome database. This database integrates existing public data resources with high-precision assembly outcomes, offering researchers a more comprehensive and accurate reference for organelle genomes.

### The annotation of chloroplast and mitochondrial genome

The chloroplast and mitochondria genome sequences were annotated with GeSEq (Tillich et al., 2017) and OGDRAW (Lohse et al., 2013). GeSeq pipeline analysis was performed using the annotation packages ARAGORN (Laslett and Canback, 2004), blatN (Kent, 2002), Chloe (Zhong, 2020) and HMMER (Eddy, 2011). GeSEq is a user-friendly online service specifically designed for the annotation of mitochondrial and chloroplast genomes. This platform enables researchers to upload unannotated DNA sequences and utilizes its database of existing high-quality annotations to identify and label genes, coding sequences, and other significant genomic features.

### Calculation of alignment length coverage ratio(ALCR)

Following the construction of local organelle genome database, all contigs assembled by the Flye software were aligned against this database using Minimap2 (Li, 2018). Subsequent to the alignment process, the next step involves calculating the Alignment Length Coverage Ratio (ALCR) for each contig. Unlike previous conventional methods, the calculation of ALCR does not solely rely on the single longest alignment region. Instead, it aggregates the lengths of all regions on the contig that align with the organelle genome database. This approach offers a more comprehensive reflection of the coverage extent between the contig and the organelle genomes. Finally, by comparing the aggregated alignment length of each contig against its total length, the alignment length coverage ratio for each contig is computed.

### Calculation of sequencing depth ratio(SDR)

The sequencing depth ratio (SDR) refers to the ratio between the sequencing depth of each contig and the average sequencing depth of the organelle genome. To calculate the SDR, the first step is to determine the average sequencing depth of the organelle genome. This is achieved by aligning the sequencing reads to the organelle genome assembled by GetOrganelle using Bowtie2 (Langmead and Salzberg, 2012), which generates a SAM file. This SAM file is then processed by Samtools (Li et al., 2009) to produce a sorted BAM file with depth information. Finally, Bedtools (Quinlan and Hall, 2010) is utilized to analyze this depth data and calculate the average sequencing depth across the organelle genome. The method for calculating the sequencing depth of each contig is identical to that used for the organelle genome. Upon completion of these calculations, the sequencing depth for each contig is divided by the average sequencing depth of the organelle genome to obtain the Sequencing Depth Ratio (SDR) for each contig.

### Identification of organelle sequences

After calculating the ALCR (Assembly Length Coverage Ratio) and SDR (Sequencing Depth Ratio) values for each contig using the locally constructed organelle genome database and second-generation sequencing data, contigs originating from the organelle genome are identified from genome assembly contigs based on ALCR and SDR filtering thresholds inputted by the user. To determine more accurate filtering thresholds, the ALCR and SDR visualization scatter plot generated after running Chlomito can be used. By utilizing this new filtering threshold, the genome assembly contigs can be filtered and selected again, resulting in more precise outcomes.

### Chromosomal-level genome assembly

The size of the genome from second-generation sequencing data was calculated using jellyfish (Marcais and Kingsford, 2011) and GenomeScope (Vurture et al., 2017) and input into Flye for contig-level genome assembly. Post assembly, contigs were corrected with racon and pilon, and redundancy was reduced using purge_dups (Guan et al., 2020). Hi-C sequencing data were aligned to the deduplicated contigs using HiC-Pro (Servant et al., 2015), and finally, Allhic (Zhang et al., 2019) was used to cluster, order, and orient the contigs based on Hi-C alignment results, achieving the final chromosomal-level genome assembly. Gaps or missing regions may be present in genome assemblies due to limitations and complexities of sequencing technologies. To obtain more complete and accurate genome sequences, we applied two approaches - Abyss Sealer (Jackman et al., 2017) and TGS-GapCloser (Xu et al., 2020) - for closing gaps in our chromosome-level genome assemblies. TGS-GapCloser utilizes long reads from third-generation sequencing (TGS) platforms to fill gaps between contigs and extend contig ends based on overlaps between contigs and long reads. Abyss Sealer is a computational tool that seals gaps in genome assemblies by aligning Illumina short reads to contig ends and performing local assembly to derive consensus sequences for gap regions.

## Results

### Overview of Chlomito workflow

#### The construction of local organelle genome database

Chlomito combines two strategies for building a comprehensive local organelle genome database as demonstrated in step 1 of Figure 1. The initial method employs GetOrganelle to assemble mitochondrial and chloroplast genomes from second-generation sequencing data. This tool is particularly valuable for assembling precise organelle genomes of species whose organelle genomic data have not yet been reported in public databases. However, the assembly of organelle genomes from short-read sequencing data with Getorganelle can sometimes be incomplete. Therefore, to ensure the comprehensiveness and reliability of the database, the second approach plays a critical role by downloading published high-quality organelle genomes from the NCBI organelle database to complement the genomes assembled by GetOrganelle. By combining these two approaches, the constructed local organelle database will overcome the limitations of relying on a single data source, offering a broad and accurate organelle genome reference for downstream analysis.

**FIGURE 1.**
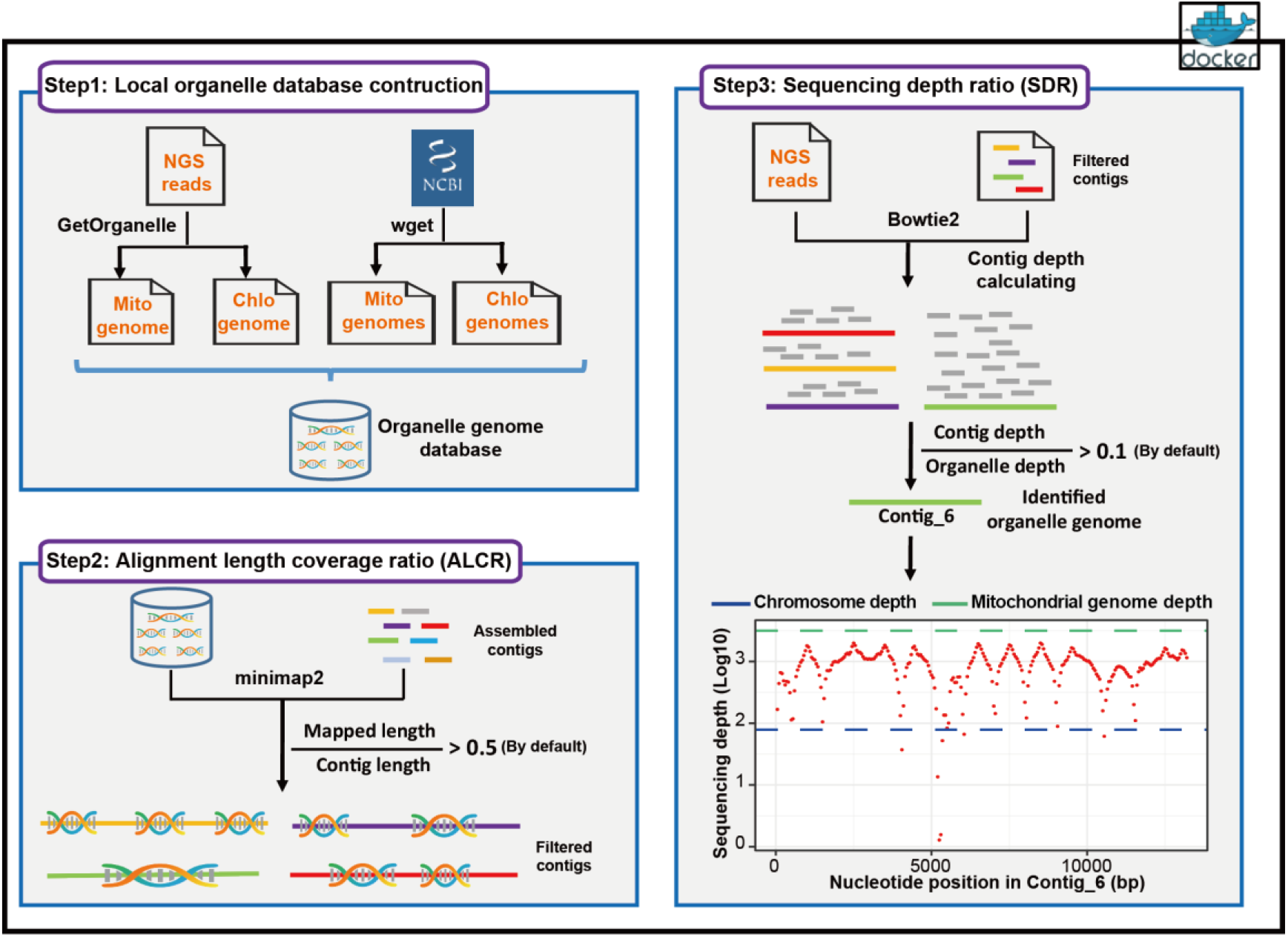
Workflow of the Chlomito software package

#### Organelle genome identification based on alignment length coverage ratio (ALCR)

After constructing the local organelle genome database, Chlomito preliminarily screens for potential organelle genomic sequences by aligning contigs assembled from third-generation sequencing (TGS) reads against this database and filtering based on the alignment results (step 2, Figure 1). The core filtering criterion at this stage was the alignment length coverage ratio (ALCR), defined as the ratio of the aligned portion of a contig to its total length. Compared to traditional methods that filter solely based on alignment length, this method can effectively distinguish original organelle sequences from those inserted into the nuclear genome via horizontal gene transfer (HGT), as HGT insertions tend to be smaller, reflected in a lower alignment length coverage ratio.

#### Organelle genome identification based on sequencing depth ratio (SDR)

To further enhance the accuracy of organelle genome sequence detection, Chlomito employs a sequencing depth ratio(SDR) method to validate the organelle genome contigs previously filtered by the ALCR criteria (step 3, Figure 1). The sequencing depth ratio(SDR) refers to the ratio of a contig’s average sequencing depth to the average sequencing depth of the organelle genome. Given that the copy number of organelle genomes is significantly higher in each cell compared to the nuclear genome, the sequencing depth ratio can be utilized to further distinguish organelle genomes from nuclear genomes. In summary, by utilizing both ALCR and SDR filtering methods, Chlomito can accurately identify organelle genomes from the total contigs. Furthermore, it can effectively reduce the misidentification of nuclear genome contigs as organelle genomes caused by horizontal gene transfer(HGT) of organelle genome.

### Overview of mitochondrial and chloroplast genomes in the NCBI database

To gain a comprehensive understanding of the characteristics of mitochondrial and chloroplast genomes across a wide range of organisms, we conducted a comparison of genome size, gene content, and other features for mitochondrial and chloroplast genomes listed in the NCBI database. The analysis revealed that the database contains 152 mitochondrial genomes and 263 chloroplast genomes derived from plants, with approximately 50% to 60% of these genomes being annotated. In comparison, the number of animal and fungal mitochondrial genomes was significantly higher, with 1,568 and 692 genomes respectively. However, the annotation rates for animal and fungal mitochondrial genomes were lower, standing at only 30%(Figure 2A). Further inspection showed that insects and ascomycetes constituted most mitochondrial genomes, while chloroplast genomes were predominantly from land plants and green algae (Figure 2B). Plant chloroplast genomes exhibited relative stability in terms of genome length and gene content, averaging around 0.15 Mb in size and containing 82 genes on average. In contrast, plant mitochondrial genomes displayed greater variability in both length and gene number as previously reported(Bendich, 2010;Oldenburg and Bendich, 2015), suggesting the potential involvement of more complex evolutionary processes(Kubo and Newton, 2008). In terms of animal mitochondrial genomes, we found a high degree of conservation, with an average size of 0.017 Mb and typically including 13 genes. Fungal mitochondrial genomes, on the other hand, had an average size of 0.063 Mb and contained an average of 14 genes (Figure 2C and 2D). These analyses not only revealed the conserved characteristics of plant chloroplast and animal mitochondrial genomes in terms of genome size and gene number, but also provided comprehensive insights into organelle genome features across taxa, laying the groundwork for further research.

**FIGURE 2.**
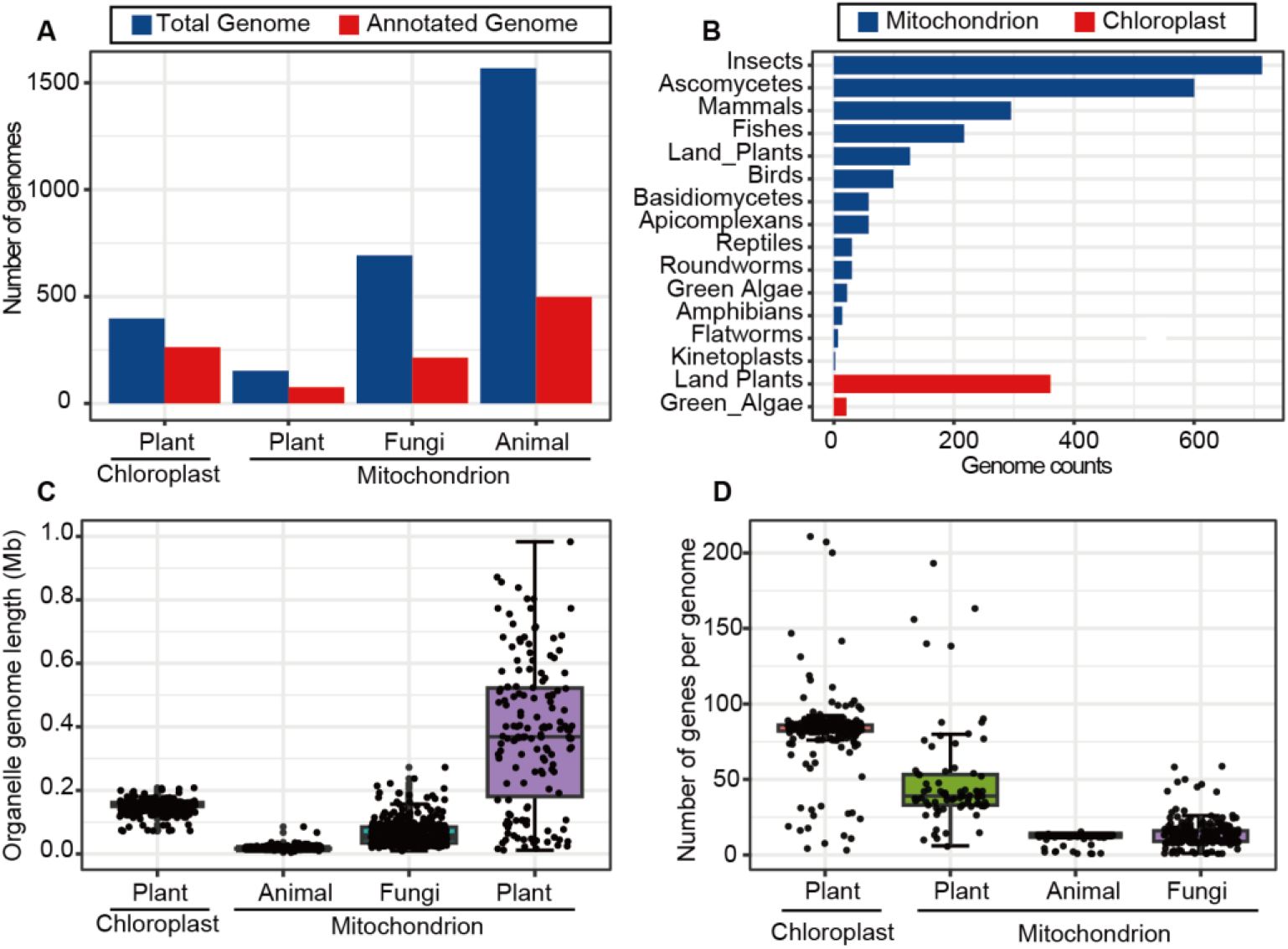
Overview of mitochondrial and chloroplast genome data in the NCBI organelle genome database. (A) Counts of total and annotated mitochondrial and chloroplast genomes across different species. (B) The number of mitochondrial and chloroplast genomes assembled for various taxonomic groups. (C) The length distribution of mitochondrial and chloroplast genomes for plants, animals and fungi. (D) The number of genes contained in mitochondrial and chloroplast genomes across different species

### The performance of Chlomito on the detection of chloroplast genome

To evaluate the performance of Chlomito in identifying chloroplast genomes, we evaluated its accuracy on publicly available sequencing data derived from Plum (Prunus salicina) and Mango (Mangifera indica). Prior to detecting chloroplast genomic sequences from all the contigs assembled from third-generation sequencing reads, we first assembled the chloroplast genomes of Mango and Plum from their NGS sequencing reads using Getorganelle, respectively. Collinearity analysis between the assembled chloroplast genomes of Mango and Plum and their published reference genomes revealed high consistency (Supplementary Figure 1), demonstrating the accuracy and reliability of the Getorganelle assemblies for downstream analysis. We then annotated the chloroplast genomes of Mango and Plum using Geseq and found that they contained similar numbers of genes with highly similar arrangements (Figure 3). To investigate the structural conservation of chloroplast genomes across diverse species, we compared the chloroplast genomes of Mango and Plum with those of other plant species including Arabidopsis thaliana and Zea mays. The results showed that chloroplast genomes were highly conserved in gene content and order across diverse plant species analyzed here (Supplementary Figure 2), implying that chloroplast genomes may be structurally conserved across diverse plants.

**FIGURE 3.**
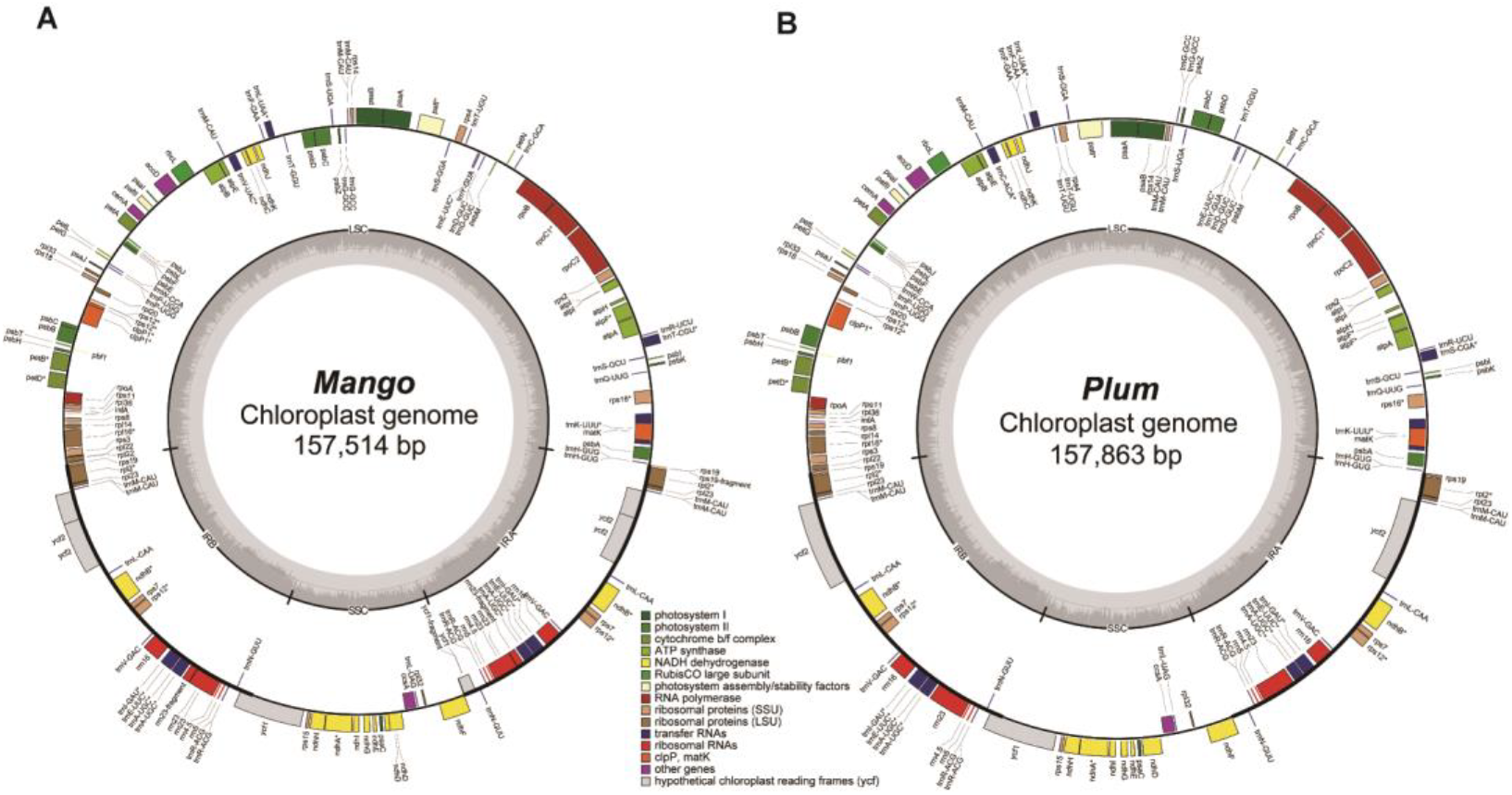
Chloroplast genome annotation of Mango(A) and Plum(B).

After constructing the local chloroplast genome databases for Mango and Plum, we employed two key metrics to identify chloroplast-derived contigs from the contigs assembled from third-generation sequencing reads. The first metric is Alignment Length Coverage Ratio (ALCR), which calculates the ratio of the aligned length of each contig to the assembled chloroplast genome versus the total length of that contig. The second metric is Sequencing Depth Ratio (SDR), which computes the sequencing depth of each contig to the average sequencing depth of the assembled chloroplast genome. Based on ALCR and SDR, we identified 3 and 5 potential chloroplast-derived contigs from Mango and Plum samples, respectively (Fig4A and 4D). These contigs showed similar alignment lengths in the GetOrganelle-assembled genomes and the NCBI chloroplast genome database (Fig4B and 4E), further validating the reliability of the chloroplast genomic sequences detected by Chlomito. Collinearity analysis revealed high consistency between these identified contigs and the chloroplast reference genomes, and the two inverted repeat regions of the chloroplast genome (IRA and IRB) were also clearly observed in the co-linearity analysis results (Fig4C and 4F). These results further confirm that these contigs were indeed derived from the chloroplast genomes and were completely detected.

Based on the alignment results of all contigs from Mango and Plum against the local chloroplast genome database, we identified 226 Mango contigs and 174 Plum contigs that aligned with the database sequences at lengths greater than 5000bp. However, the proportion of each contig’s length that aligned to the database was low, with less than 10% coverage (Supplementary Figure 3). The chloroplast genomic fragments present in these ultra-long contigs may have been inserted into the nuclear chromosomes through horizontal gene transfer from the chloroplast genome. These results indicate that the traditional method of filtering out chloroplast genomes based solely on alignment length may erroneously exclude some ultra-long nuclear contigs that contain only a small proportion of chloroplast genomic content. Therefore, adopting more refined filtering criteria, such as ALCR and SDR, is critical to accurately differentiate contigs that are truly from the organelle genome versus those inserted into the nuclear genome through HGT.

### The performance of Chlomito on the detection of mitochondrial genomes

After validating the accuracy of Chlomito for identifying chloroplast genome sequences, we further tested its ability to accurately identify mitochondrial genome sequences. Mitochondrial genomes of Mango and Plum were first assembled from next-throughput sequencing (NGS) data using the GetOrganelle software. Unlike the single complete chloroplast genomes assembled by Chlomito, the reconstructed mitochondrial genomes of Mango and Plum were composed of multiple fragments. The assembled Mango mitochondrial genome consisted of 15 sequences totaling 0.48 Mb, while the Plum mitochondrial genome was composed of 11 sequences amounting to 0.36 Mb. To validate the accuracy of the mitochondrial genomes assembled by GetOrganelle, we performed collinearity analysis between the assembled genomes and the published mitochondrial reference genomes for Mango (MZ751075.1) and Plum (OK563724.1). The results showed that the assembled mitochondrial genomes had high collinearity with the reference genomes and covered most regions of the reference genome (Supplementary Figure 4), indicating the high accuracy and completeness of the GetOrganelle-assembled mitochondrial genomes. To evaluate the conservation of structures within mitochondrial genomes across different plant species, we annotated and compared the mitochondrial genomes of Mango, Plum, Zea mays, and Arabidopsis thaliana with the OGDRAW website. The results showed that unlike chloroplast genomes, the mitochondrial genomes from these diverse species did not show conserved gene content or gene order (Supplementary Figure 5). Subsequently, we integrated the assembled mitochondrial genomes into the mitochondrial genome database downloaded from NCBI to establish a comprehensive local mitochondrial genome database.

**FIGURE 4.**
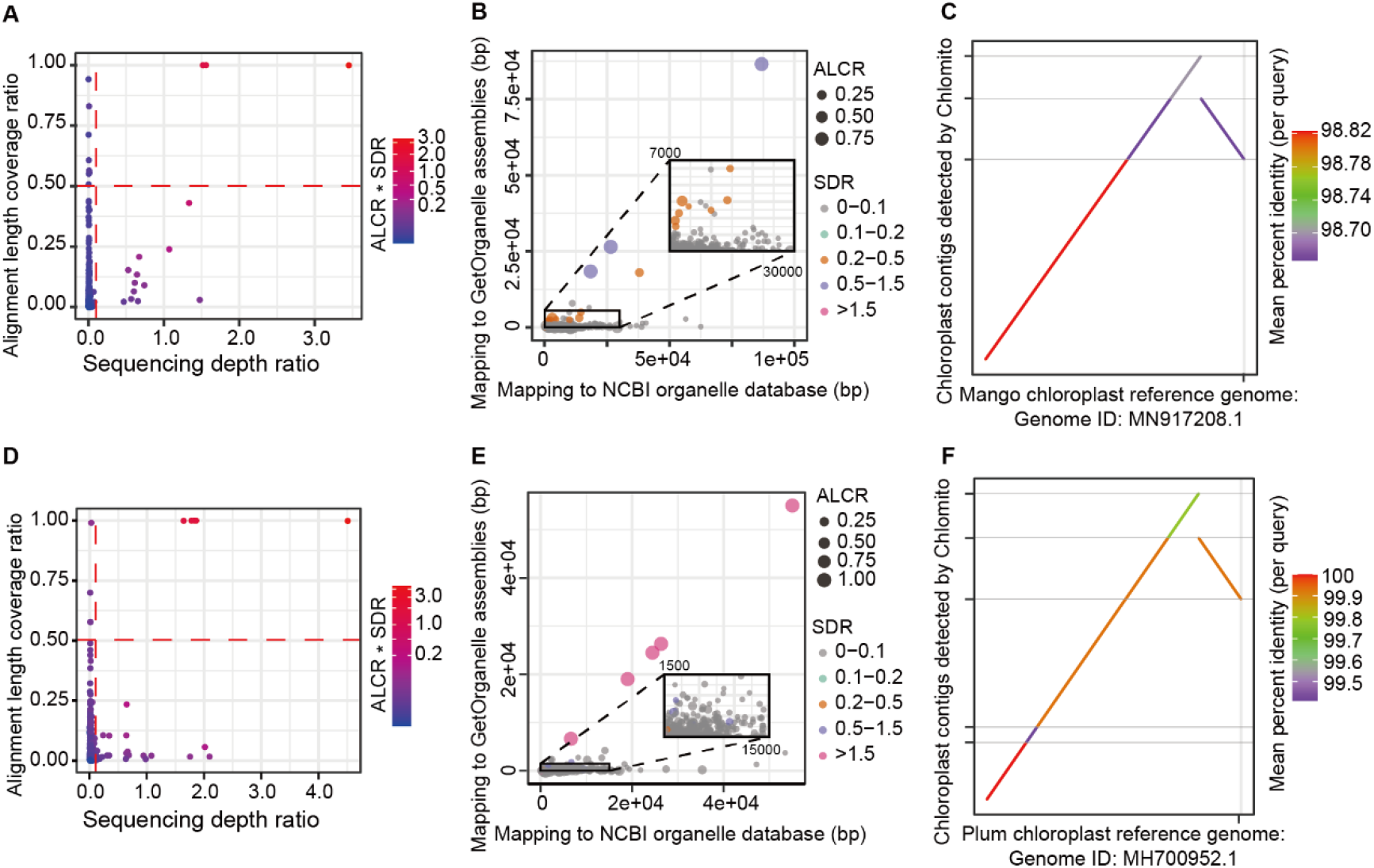
Identification and analysis of Chloroplast-derived contigs from Mango and Plum Samples. (A, D) Identification of chloroplast-derived contigs in Mango (A) and Plum (D) based on ALCR and SDR metrics. (B, E) Alignment lengths of Mango (B) and Plum (E) contigs with chloroplast genomes assembled using GetOrganelle and downloaded from NCBI database. (C, F) Collinearity analysis of Mango (C) and Plum (F) contigs identified by Chlomito against published chloroplast reference genomes.

**FIGURE 5.**
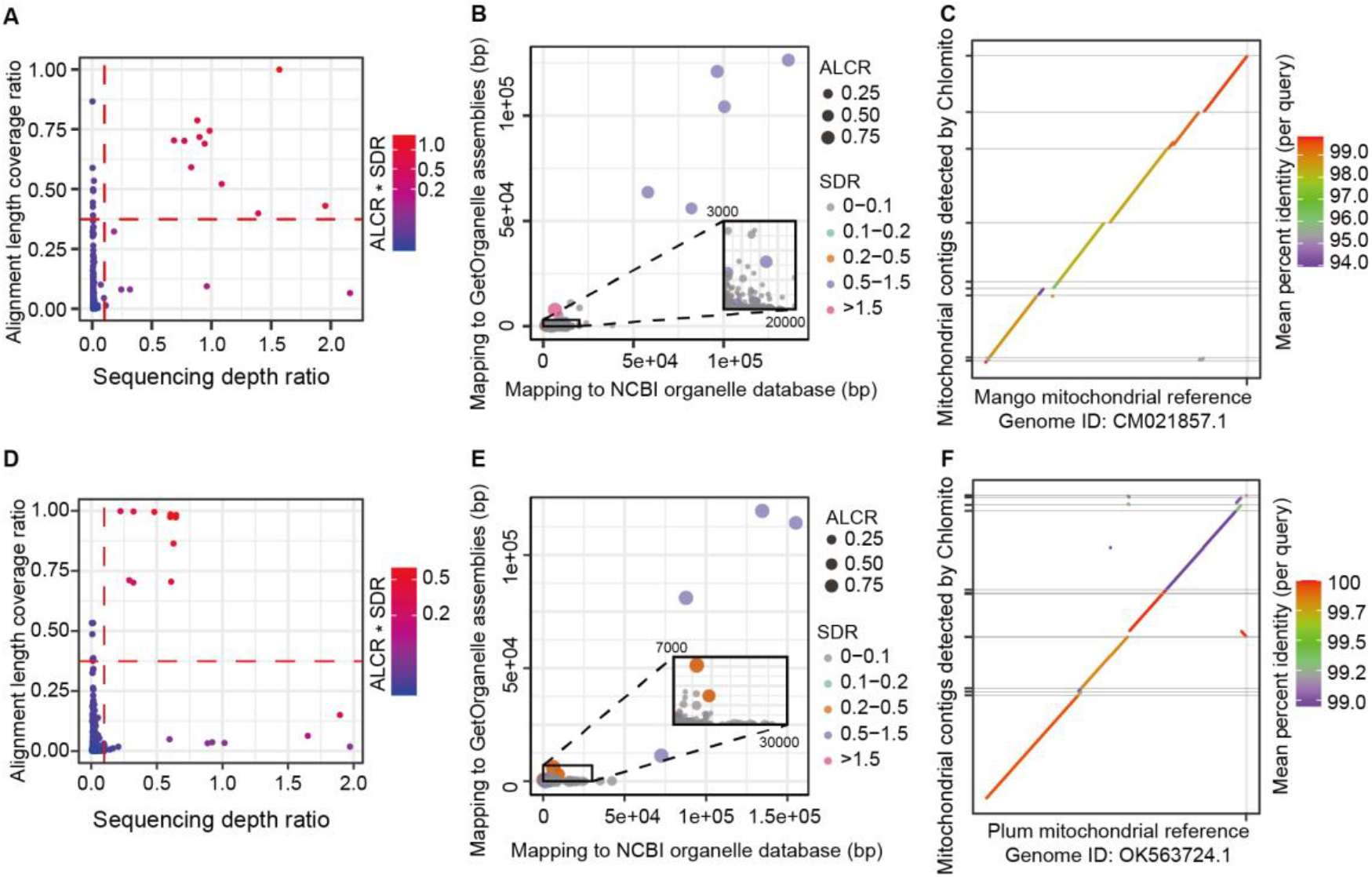
Identification and analysis of mitochondrial-derived contigs from Mango and Plum samples. (A, D) Identification of mitochondrial-derived contigs in Mango (A) and Plum (D) using ALCR and SDR metrics. (B, E) Alignment lengths of contigs with GetOrganelle-assembled and NCBI-derived mitochondrial genomes for Mango (B) and Plum(E). (C, F) Collinearity analysis of Chlomito-identified contigs against published mitochondrial reference genomes for Mango (C) and Plum (F).

After constructing local mitochondrial genome databases for Mango and Plum, we utilized these databases along with second-generation sequencing data to calculate the Alignment Length Coverage Ratio (ALCR) and Sequencing Depth Ratio (SDR) for each contig. Based on the ALCR and SDR parameters, we identified 11 and 10 contigs likely originating from the mitochondrial genome in Mango and Plum, respectively (Fig5A and 5D). These contigs also exhibited similar alignment lengths in the mitochondrial genomes assembled by GetOrganelle and the NCBI mitochondrial genome database (Fig5B and 5E). Moreover, Collinearity analysis revealed a high consistency between the identified contigs and the mitochondrial reference genomes of Mango and Plum, with the majority of the mitochondrial genome regions being covered by these contigs (Fig5C and 5F). Similar to findings in chloroplast genome identification, we also detected 116 and 52 contigs with alignment lengths exceeding 5000bp but exhibiting low coverage (less than 10%) in Plum and Mango, respectively (Supplementary Figure 6). These results indicate that large fragment horizontal gene transfer exists in both mitochondrial and chloroplast genomes. Therefore, more stringent ALCR and SDR filtering criteria are essential to replace the traditional organelle genome identification methods that rely solely on alignment length.

**FIGURE 6.**
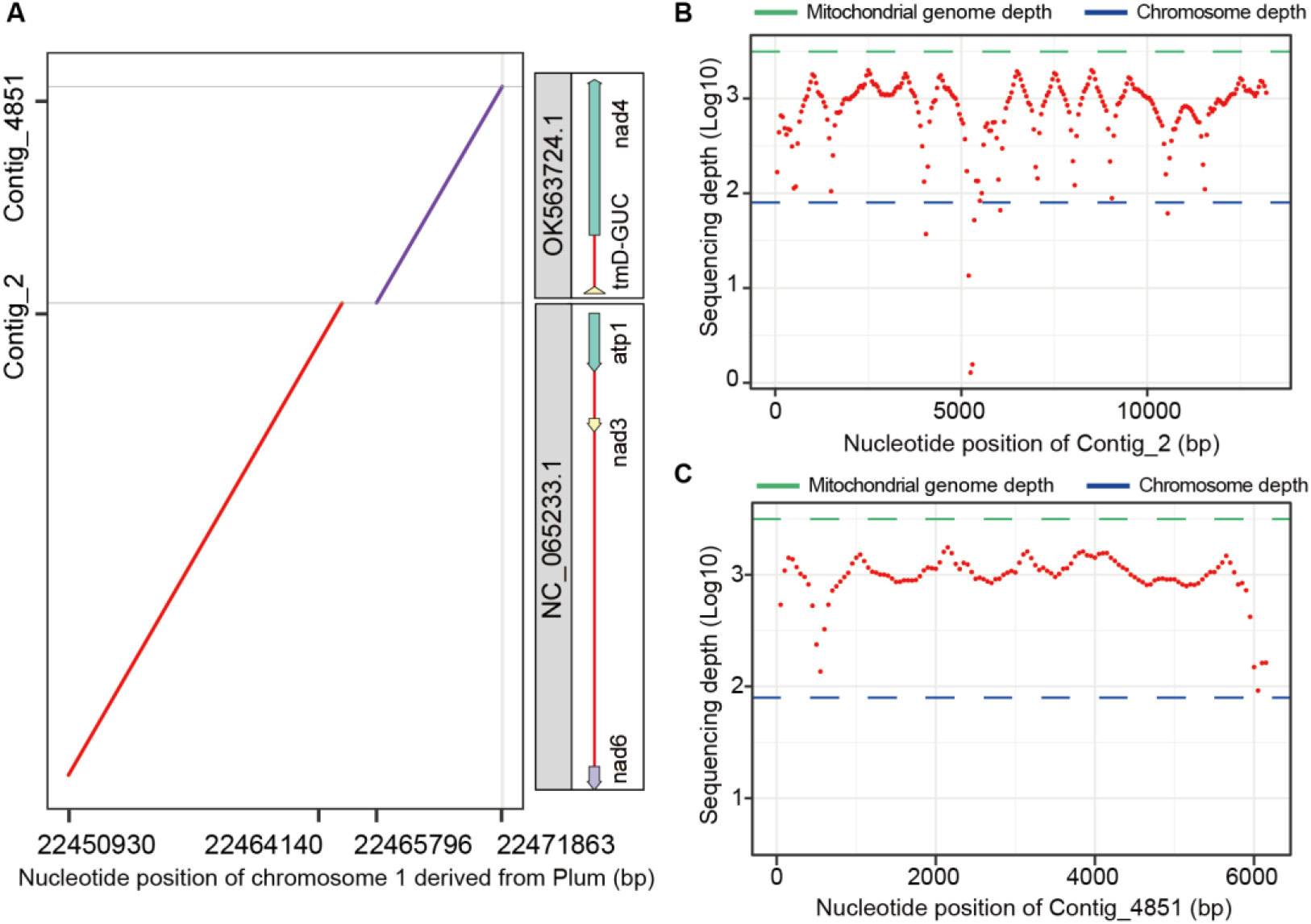
Analysis of organelle genome contamination in chromosome assembly of Plum. (A)Sequence alignment of two mitochondrial-derived contigs (identified by Chlomito) with assembled chromosome 1 of Plum. (B, C) Sequencing depth analysis for contig_2 (B) and contig_4851 (C), comparing to the average depth of chromosome and mitochondrial genomes.

### Organelle genome contamination in chromosome assembly

To evaluate the impact of organelle genome sequences on the accuracy of chromosome assembly, we aligned the chromosomes assembled from all contigs without filtering organelle sequences to the contigs identified as organelle DNA by Chlomito. The aligned result revealed that two mitochondrial genome fragments (contig_2 and contig_4851) identified by Chlomito were erroneously assembled into chromosome 1 of Plum (Figure 6A). These fragments showed perfect matches with mitochondrial reference genome (NC_065233.1 and OK563724.1) in the NCBI organelle genome database. Further analysis of sequencing depth demonstrated that the coverage of these two contigs was much higher than the average depth of the chromosomal genome, and was close to the average depth of the organelle genome (Fig. 6B and 6C). This is consistent with the characteristic that organelle genomes have higher copy numbers than nuclear genomes. These results confirm that the contigs integrated into the chromosome were indeed derived from the mitochondrial genome. Furthermore, these findings highlight that unfiltered organelle sequences can truely contaminate the nuclear genome during chromosome-level genome assembly. Therefore, the preliminary identification and exclusion of organelle genome sequences using tools like Chlomito are curcial for ensuring the accuracy and integrity of chromosome assembly.

## Discussion

In this study, we have developed a software called Chlomito that provides an innovative approach for accurately identifying organelle genome sequences from complex genomic assemblies. This method significantly improves the accuracy of recognizing organelle genomic fragments by integrally applying two metrics. The first metric, Alignment Length Coverage Ratio (ALCR), is calculated as the ratio of the aligned length of each contig to the assembled organelle genome versus the total length of that contig. The introduction of ALCR will significantly reduce the likelihood of incorrectly identifying nuclear genomic sequences as organelle genome sequences, especially for those organelle genome sequences that may have been inserted into the nuclear genome via HGT. The second metric, Sequencing Depth Ratio (SDR), which compares the sequencing depth of each contig to the average sequencing depth of the organelle genome. The application of SDR will provides an additional robust filtering dimension, further ensuring the identification of sequences truly originating from organelle genomes among all the assembled contigs. Compared to conventional methods only relying on reference alignment length filtering, our approach effectively overcomes the challenge posed by horizontal gene transfer (HGT) events that make it difficult to distinguish organelle genomes from complex genomic assemblies.

To evaluate the accuracy of Chlomito in detecting organelle genome sequences, we tested it using sequencing data from Mango (Mangifera indica) and Plum (Prunus salicina). By performing collinearity analysis between the contigs detected by Chlomito as being from the organelle genomes and the organelle reference genomes in the NCBI database, we observed high collinearity between these contigs and the organelle reference genomes. This demonstrates that the contigs detected by Chlomito are indeed from the organelle genomes. More importantly, the contigs detected by Chlomito were able to cover the vast majority of the organelle reference genome regions, which not only verified its high accuracy in the detection of organelle genome sequences but also demonstrated its capability in effectively identifying the entire organelle genome sequence. These results confirm the efficacy of Chlomito as a tool for identifying plant organelle genomes, providing reliable technical support for subsequent studies in organelle genomics.

Furthermore, we noted that in addition to the contigs explicitly identified as organelle genomes by Chlomito, there were more than a hundred contigs with alignment lengths to the reference organelle genomes exceeding 5000bp but exhibiting low coverage (less than 10%) relative to the length of the contigs themselves. This phenomenon may be attributable to the insertion of organelle genomes into the nuclear genome via horizontal gene transfer (HGT) events. HGT is a significant mechanism in the evolutionary process, particularly in the exchange of genetic information between organelle genomes and host nuclear genomes. Our findings provide further evidence that HGT may be widespread across different species and plays a crucial role in the evolutionary interaction between organelle and nuclear genomes. In this context, it is particularly important to accurately distinguish sequence exchanges between organelle and nuclear genomes caused by HGT. Chlomito is designed to address this challenge by employing two filtering criteria - ALCR and SDR - to effectively differentiate contigs that are truly from the organelle genome versus those inserted into the nuclear genome through HGT. The application of this strategy not only enhances the accuracy of organelle genome identification but also deepens our understanding of the complex interactions between organelle and nuclear genomes.

Additionally, we explored the impact of organelle genome sequences on the accuracy of chromosomal-level genome assembly. The results revealed that without the removal of organelle genome sequences, segments of organelle genomes may be erroneously assembled into chromosomes by genome assembly software, compromising the accuracy of the chromosomal assembly. These results clearly demonstrate the critical importance of identifying and eliminating organelle genome contamination prior to chromosomal-level assembly to ensure the fidelity of the assembly outcomes. Therefore, the development of tools like Chlomito is critical for improving the accuracy of chromosomal-level genome assembly, and enhancing the quality and reliability of genome assembly outcomes in scientific research.

In summary, the development of Chlomito provides an accurate and effective solution for identifying and removing organelle DNA fragments from genome assembly contigs, holding significant value in improving the quality of chromosome assembly and deepening our understanding of the complex interactions between organelle and nuclear genomes.

## Funding

The work was supported by the General Program of National Natural Science Foundation of China (31970622).

## Author contributions

Conceptualization, W.S., H.J. and W.P.; the development of Chlomito program and data analysis, W.S., C.L. and Y.J.; writing—draft preparation, W.S. and Y.L.; writing— review and editing, W.P., D.S., Y.H. and H.J.; supervision, H.J., W.P. and Y.H.. All authors have read and agreed to the published version of the manuscript.

## Competing interests

The authors declare no conflict of interest.

## Supplementary Materials

**Figure S1.**
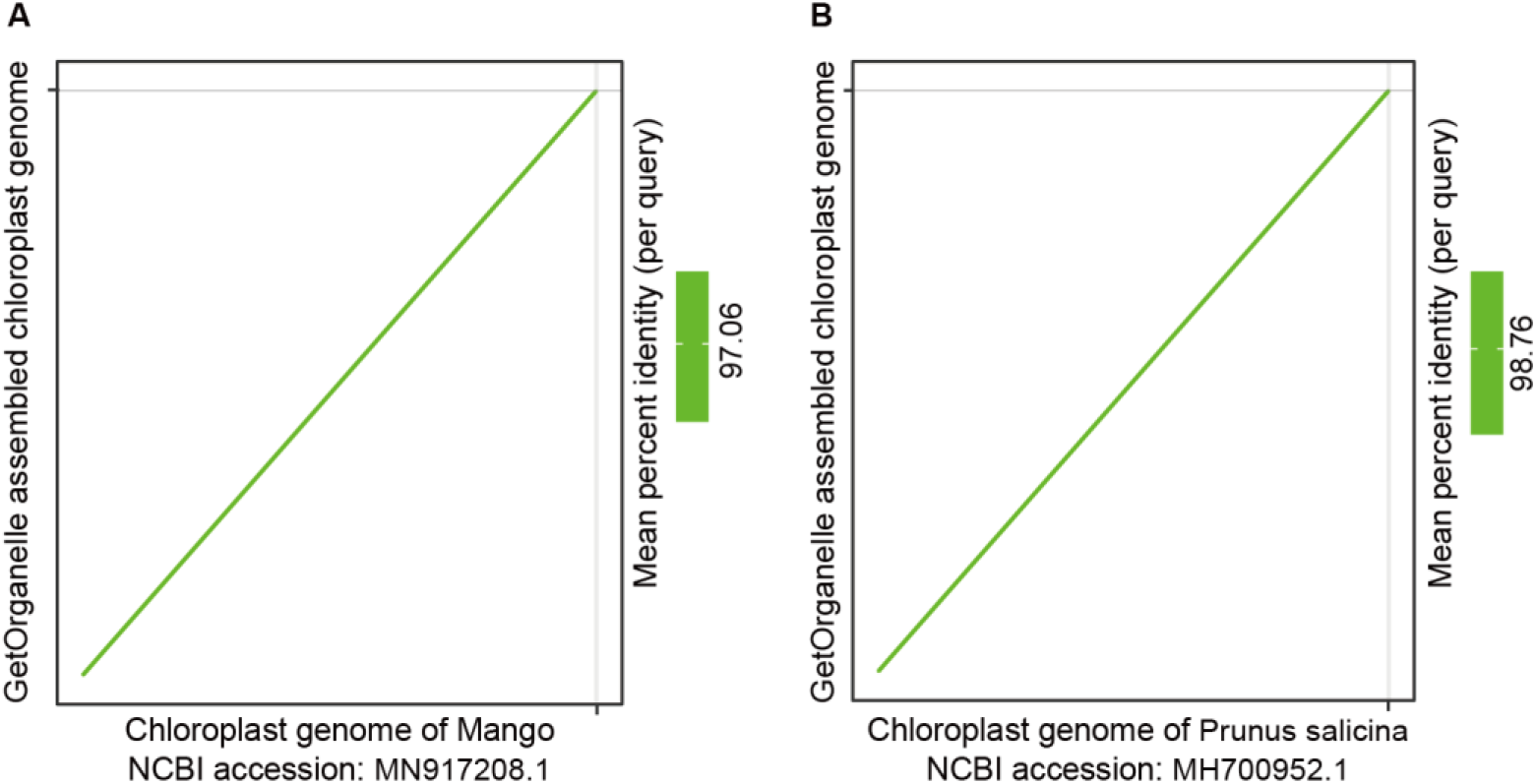
Collinearity comparison of chloroplast genomes of Mango (A) and Plum (B) assembled by GetOrganelle with chloroplast reference genomes from NCBI.

**Figure S2.**
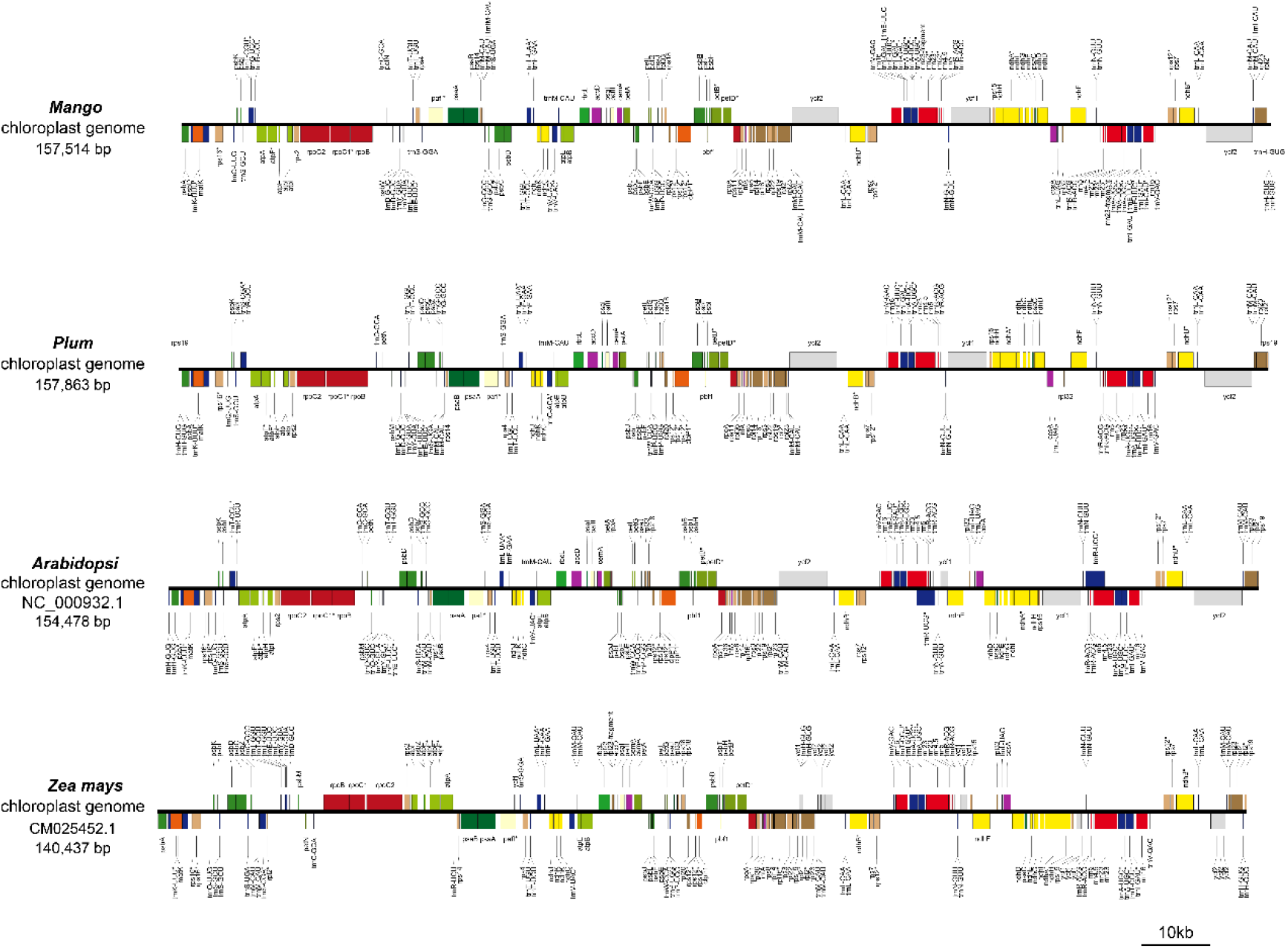
Annotation and comparison of chloroplast genomes of Mango, Plum, Arabidopsis, and Zea mays.

**Figure S3.**
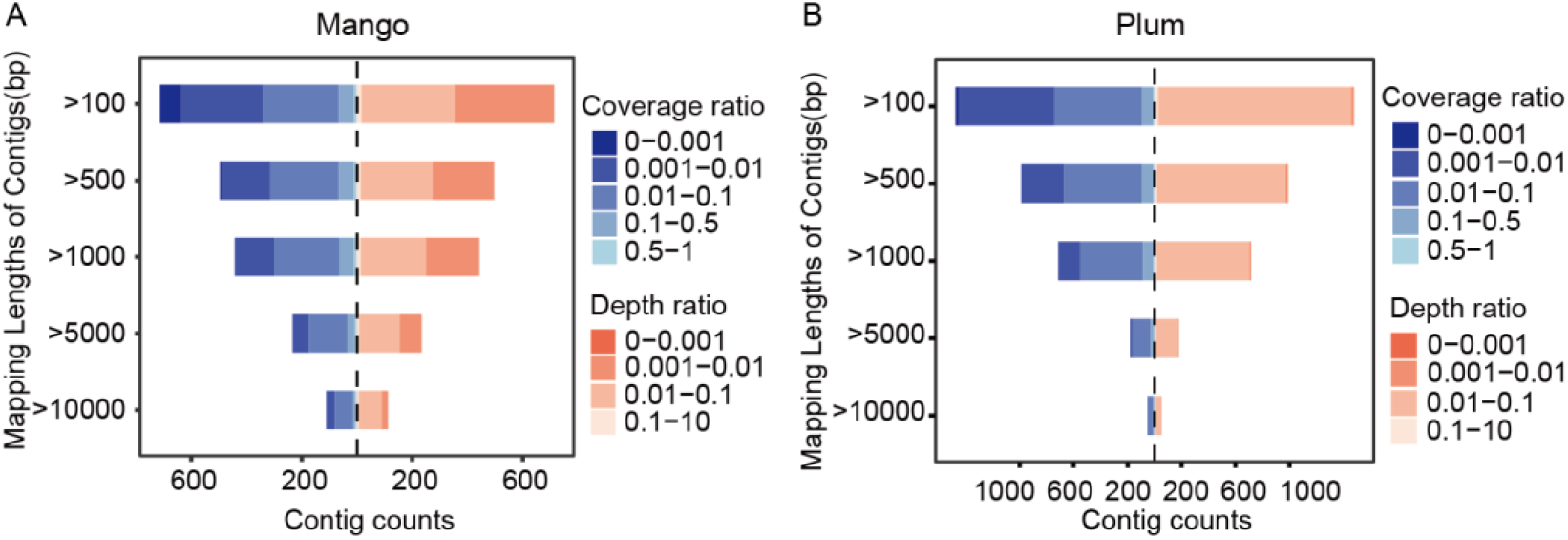
Comprehensive analysis of alignment length coverage ratio, sequencing depth ratio and contig alignment length with chloroplast reference genomes in Mango(A) and Plum(B) Samples.

**Figure S4.**
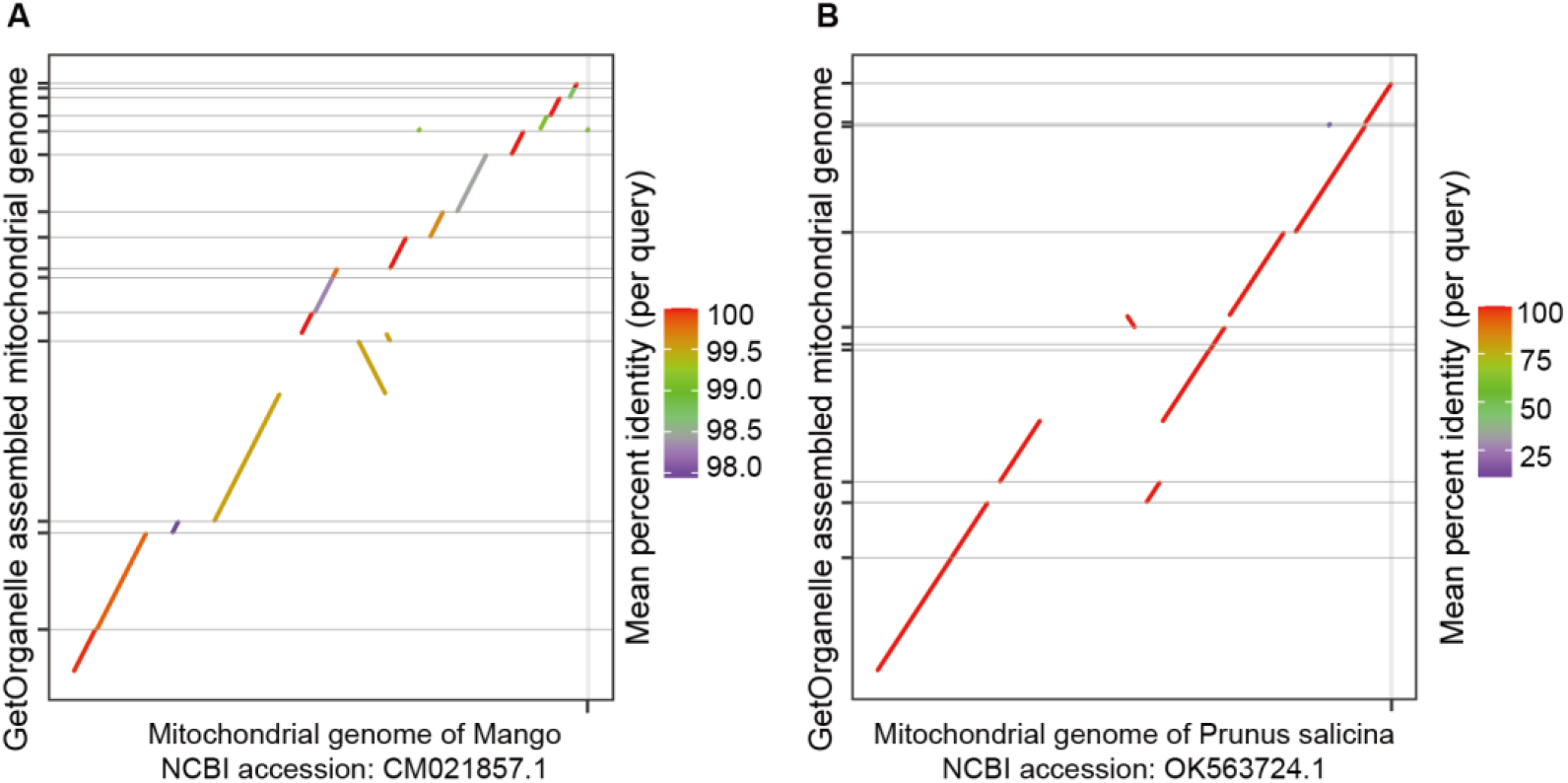
Collinearity comparison of mitochondrial genomes of Mango (A) and Plum (B) assembled by GetOrganelle with mitochondrial reference genomes from NCBI.

**Figure S5.**
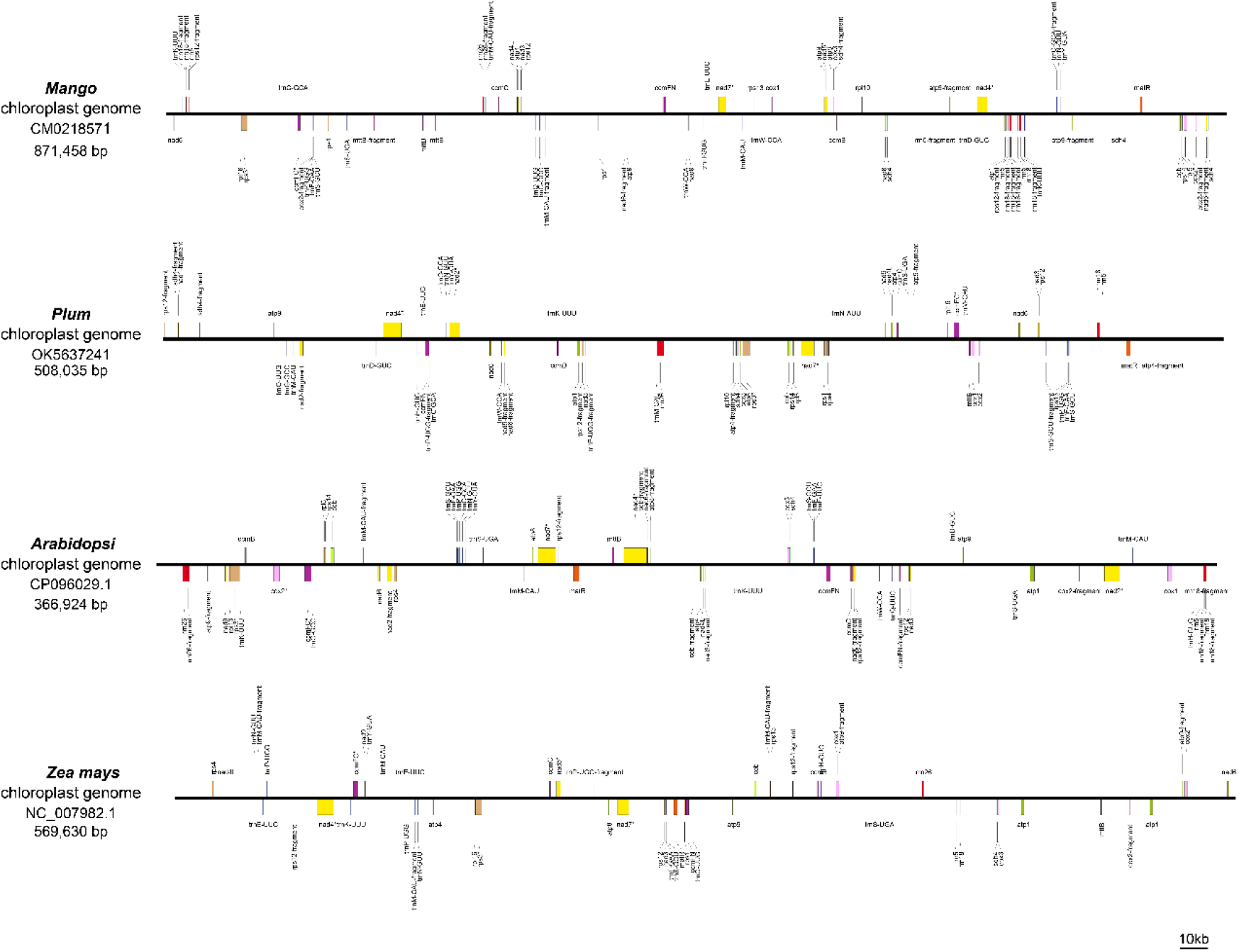
Annotation and comparison of mitochondrial genomes of Mango, Plum, Arabidopsis, and Zea mays.

**Figure S6.**
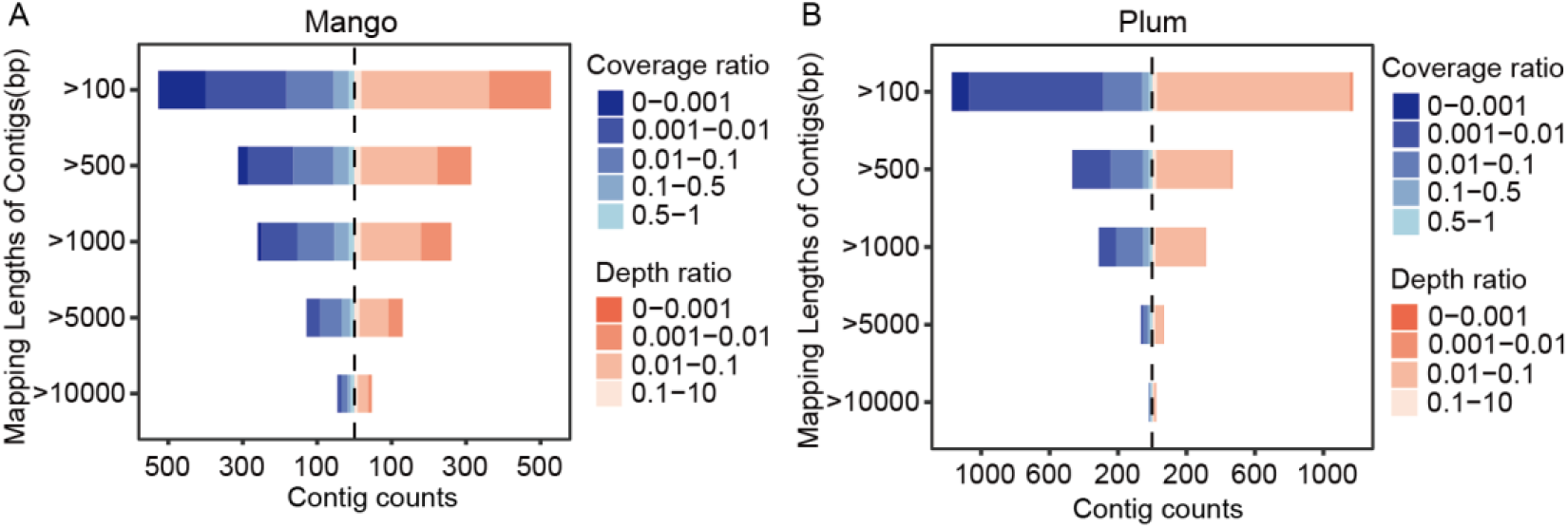
Comprehensive analysis of alignment length coverage ratio, sequencing depth ratio and contig alignment length with mitochondrial reference genomes in Mango(A) and Plum(B) Samples.

## Notes

### Competing Interest Statement

The authors have declared no competing interest.

